# TREEasy: an automated workflow to infer gene trees, species trees, and phylogenetic networks from multilocus data

**DOI:** 10.1101/706390

**Authors:** Yafei Mao, Siqing Hou, Evan P. Economo

## Abstract

Multilocus genomic datasets can be used to infer a rich set of information about the evolutionary history of a lineage, including gene trees, species trees, and phylogenetic networks. However, user-friendly tools to run such integrated analyses are lacking, and workflows often require tedious reformatting and handling time to shepherd data through a series of individual programs. Here, we present a tool written in Python—TREEasy—that performs automated sequence alignment (with MAFFT), gene tree inference (with IQ-Tree), species inference from concatenated data (with IQ-Tree), species tree inference from gene trees (with ASTRAL, MP-EST, and STELLS2), and phylogenetic network inference (with SNaQ and PhyloNet). The tool only requires FASTA files and nine parameters as inputs. The Tool can be run as command line or through a Graphical User Interface (GUI). As examples, we reproduced a recent analysis of staghorn coral evolution, and performed a new analysis on the evolution of the WGD clade of yeast. The latter revealed novel inferences that were not identified by previous analyses. TREEasy represents a reliable and simple tool to accelerate research in systematic biology (https://github.com/MaoYafei/TREEasy).

## Introduction

The inference of evolutionary history from molecular data is a core goal of modern evolutionary biology (Barraclough and Nee, 2001; Soltis and Soltis, 2018). With the increasing availability of large-scale multilocus datasets and advances in computational power, phylogenetic methods have diversified in the past two decades (Delsuc et al., 2005; Liu et al., 2015b). Instead of inferring a bifurcating tree with a single locus as the focus of analysis, biologists regularly infer populations of trees representing the histories of different loci (Edwards et al., 2007; Gadagkar et al., 2005). From these, species tree methods can be used which take into account the fact that gene trees can be discordant with species trees even under a bifurcating evolutionary history (Kubatko and Degnan, 2007; Lambert et al., 2015; Page and Charleston, 1997; Shen et al., 2016; Tonini et al., 2015). In addition, evolutionary histories are not always bifurcating (Bravo et al., 2019; Degnan and Rosenberg, 2009; Gadagkar et al., 2005; Page and Charleston, 1997), introgression is relatively common occurrence across the tree of life (Berner and Salzburger, 2015; Bravo et al., 2019; Morrison, 2014; Xu, 2000). Phylogenetic network methods can be used to infer evolutionary histories that include reticulation (Bastide et al., 2018; Huson and Bryant, 2005).

In total, these methods increasingly reflect the complexity of evolution, and for each of these analyses types, multiple programs are available to the researcher. For example, methods allowing inference of species trees and phylogenetic networks include: NJst (Liu and Yu, 2011), MP-EST (Liu et al., 2010), ASTRAL (Mirarab et al., 2014), STELLS2 (Pei and Wu, 2017), Guenomu (de Oliveira Martins and Posada, 2017), SNaQ (Solís-Lemus et al., 2017) and PhyloNet (Wen et al., 2018). However, each method requires gene tree input and control files in different format. In particular, ASTRAL requires an unrooted gene tree list whereas MP-EST requires a rooted gene tree list. In addition, Guenomu, a Bayesian hierarchical model, requires posterior distributions of gene trees. SNaQ runs in Julia language whereas PhyloNet runs in a command line with a special control file. Thus, there is much preset work to run these tools. STRAW was developed as a Web-based server requiring a gene tree list as input to infer species tree with STAR, MP-EST, and NJst (Shaw et al., 2013), but it only runs for rooted gene trees and cannot infer phylogenetic networks.

Together, there is no tool to integrate sequence alignment, gene tree reconstruction, species tree, and phylogenetic network inferences into a single run. This means the researcher has to spend considerable time to figure out how to reformat files and run each individual program, reducing efficiency and making it less likely that a broad range of methods and programs are used. To address this problem, we present a multi-thread open-source tool, named TREEasy, to shepherd data through a series of programs to infer gene trees, species trees, and phylogenetic networks from molecular sequences.

## TREEasy architecture

TREEasy is written in Python integrating sequence alignment, gene tree reconstruction, species tree inference, and phylogenetic network inference. BioPython must be installed and a few executable dependencies are needed: MAFFT (Nakamura et al., 2018), Translatorx (Abascal et al., 2010), AMAS (Borowiec, 2016), IQ-TREE (Nguyen et al., 2014), ASTRAL (Mirarab et al., 2014), MP-EST (Liu et al., 2010), STELLS2 (Pei and Wu, 2017), PhyloNet (Wen et al., 2018) and SNaQ (Solís-Lemus et al., 2017). Molecular sequences in FASTA format (SNP, microsatellites, protein-coding sequences, etc.) are mandatory inputs. In addition, to identify multiple individuals in a species and identify mismatches between species names in different gene files, two text files including species numbers and gene names respectively are needed. There are 3 subroutines in TREEasy as follows (Figure 1).

**Figure 1.**
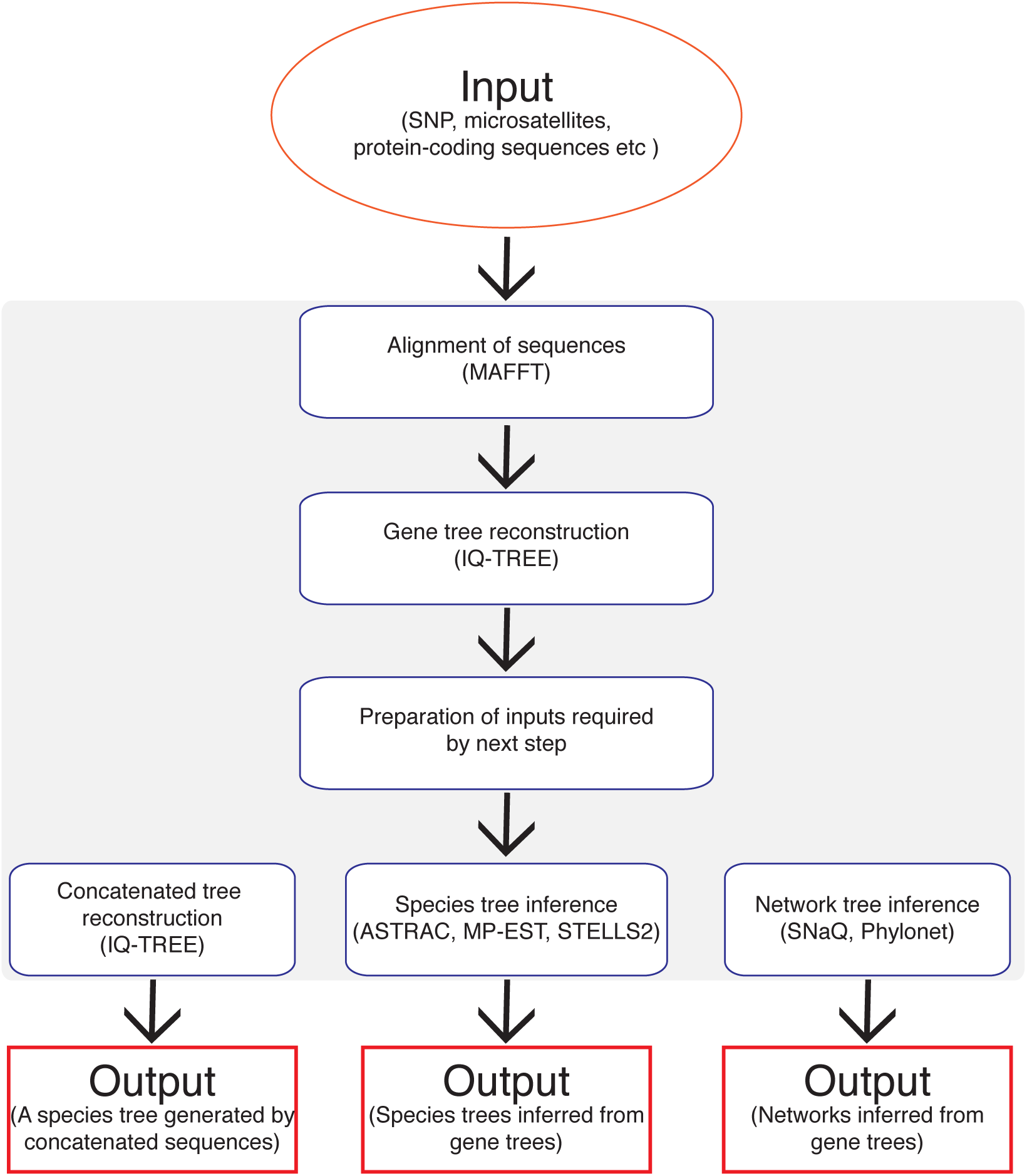
Workflow in TREEasy. The orange oval represents inputs. Blue boxes represent subroutines and red boxes represent outputs.

### (1) Gene tree reconstruction

Molecular sequences are aligned using MAFFT with localpair model and then gene trees are reconstructed with Maximum likelihood (ML) method in IQ-TREE with model selection. This process runs as parallel with the threading module in python.

### (2) Species tree inference

Firstly, alignments of multi-loci are concatenated to build a concatenated species tree using IQ-TREE. Then, the tool selects gene trees of which bootstrap values are greater than B (B is a preset parameter from 0 to 100) in order to avoid uncertainty of gene tree reconstruction. Next, un-rooted selected gene trees generated by IQ-TREE are put together as input to infer a species tree using ASTRAL. Meanwhile, the un-rooted gene trees are rooted with a preset parameter R (species name(s)) and then the rooted gene trees are used to infer species trees using STELLS2 and MP-EST.

### (3) Phylogenetic network inference

A species tree generated by ASTRAL and the un-rooted gene trees are used to infer a phylogenetic network using SNaQ. Then, the rooted gene trees are used to infer a phylogenetic network using PhyloNet.

The tool can be run as command line or through Graphical User Interface (GUI) for users. The GUI interface can be seen in Figure 2.

**Figure 2.**
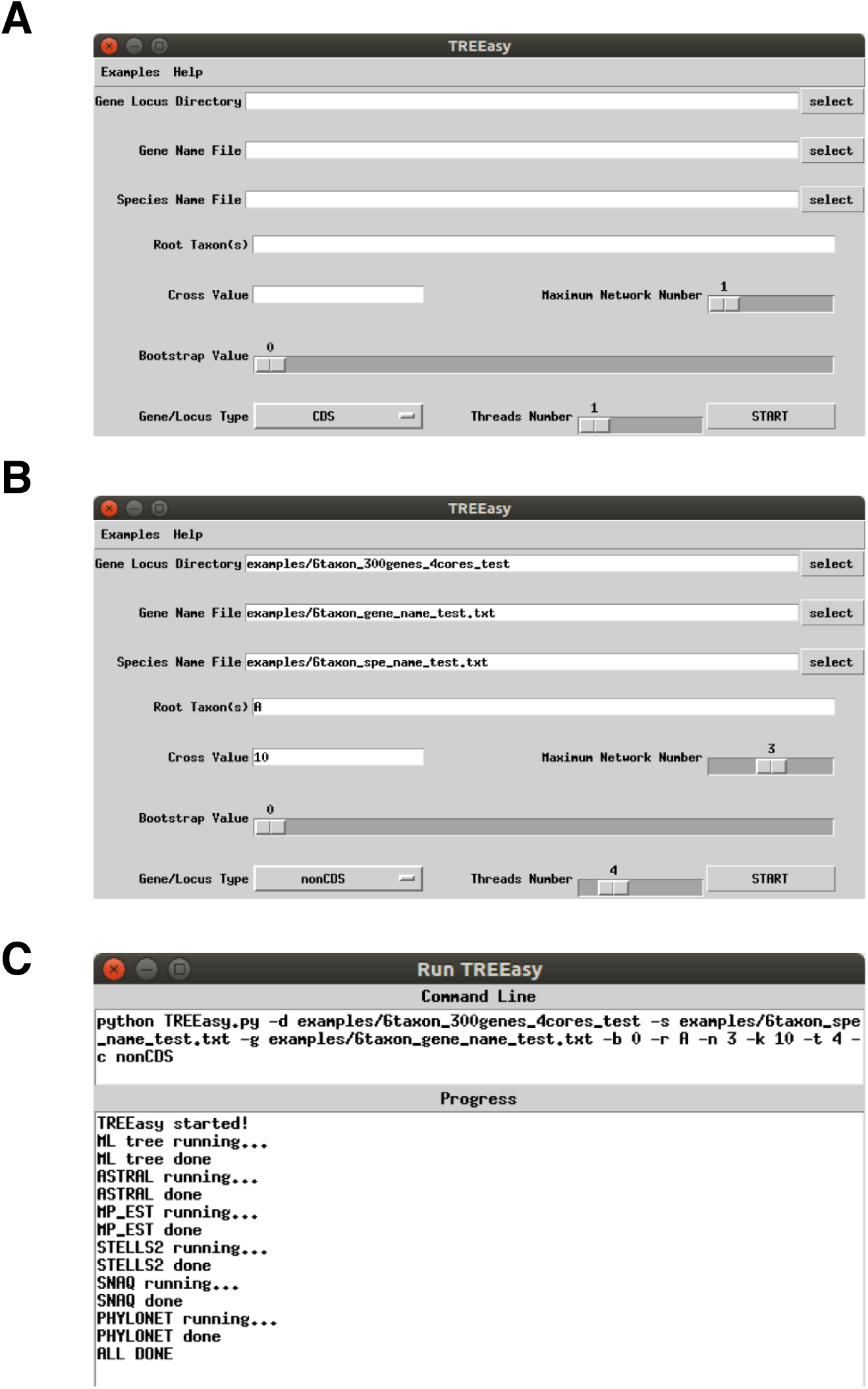
GUI windows of TREEasy. (A) The start window. (B) The window shows the simulated data example. (C) The window shows a successful run of the simulated data example.

## Results

### Evaluation with simulated data

To evaluate the performance of TREEasy, we used simulated data from the previous study (Solís-Lemus et al., 2017) to evaluate following aspects of TREEasy (Figure 3): (1) running time and memory usage with different processors; (2) with different gene numbers; (3) with different taxon numbers.

**Figure 3.**
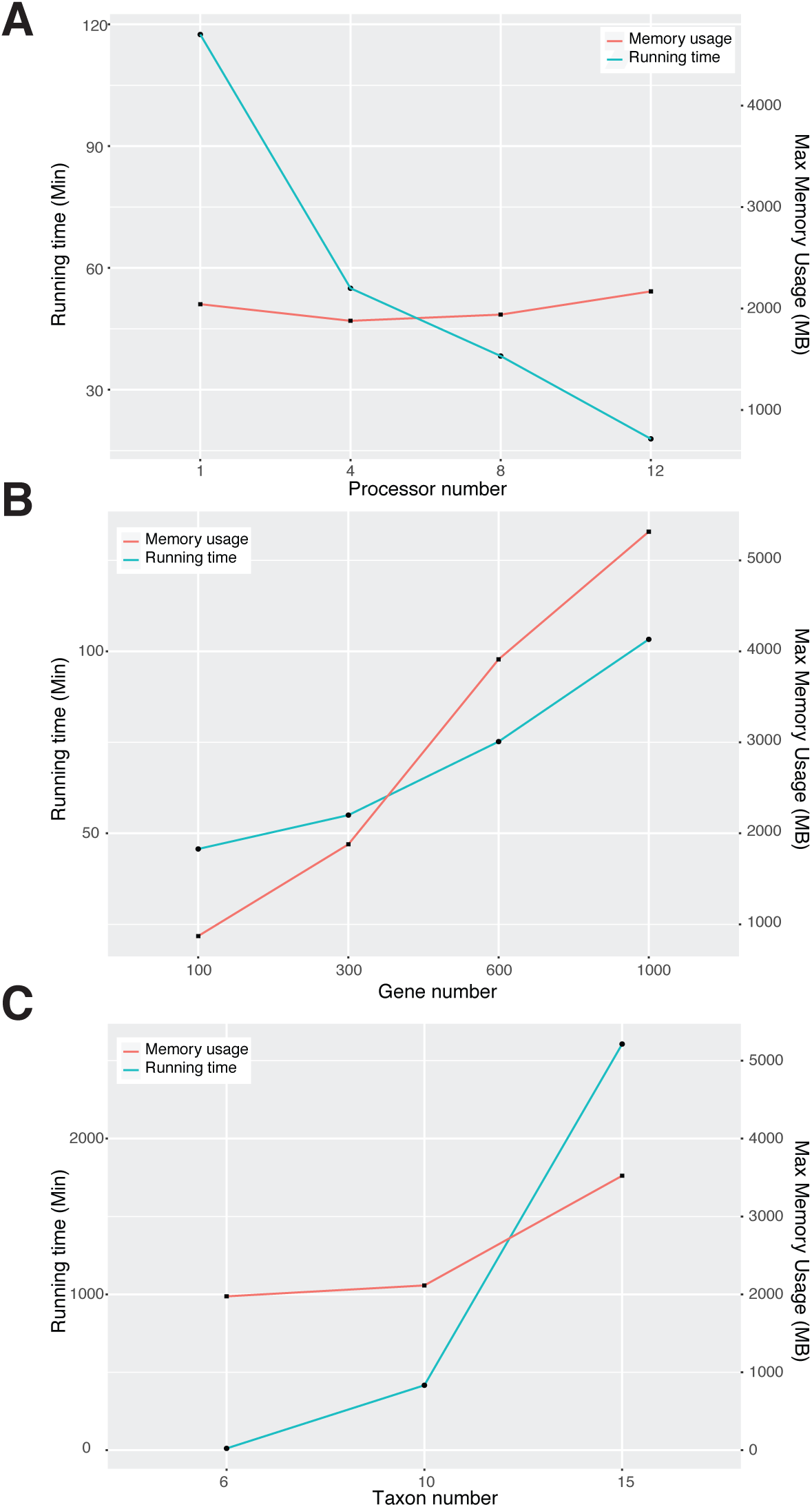
Estimation of TREEasy on simulated data. (A) Running time and maximum memory usage versus processor number. (B) Running time and maximum memory usage versus gene number. (C) Running time and maximum memory usage versus taxon number.

First, with 6 taxa and 300 genes, we found that the running time decreased with the increase of processor number and the speed with 12 processors was 6.5 times faster compared to with 1 processor. Yet, the maximum memory usage did not show a significant change. Second, with 6 taxa and 4 processors, we found that the running time and maximum memory usage increased with increase of gene number. Third, with 300 genes and 12 processors, we found that the running time excluding the run of PhyloNet increased from 6 taxa to 15 taxa, but the maximum memory usage was increased dramatically from 10 taxa to 15 taxa. It is worth noting that PhyloNet running was extremely slow with > 10 taxa (> 7 days) and thus we excluded the running of PhyloNet in this analysis.

### Empirical data validation

As a second test, we used TREEasy to reproduce a previous analysis of *Acropora* genome evolution that inferred reticulation events among five coral species (Mao et al., 2018). The *Acropora* data included 4,945 single-copy orthologs among five *Acropora* species. The whole process took ∼13 hours with maximum memory usage: 3,992 Mb, running on 8 processors. We found that the concatenated species tree has the same topology as the other species trees inferred from gene trees with ASTRAL, MP-EST, and STELLS2. Then, the inferred phylogenetic network topologies with SNaQ and PhyloNet are identical (Figure 4). Both of these results are coincident with our previous study (Mao et al., 2018).

**Figure 4.**
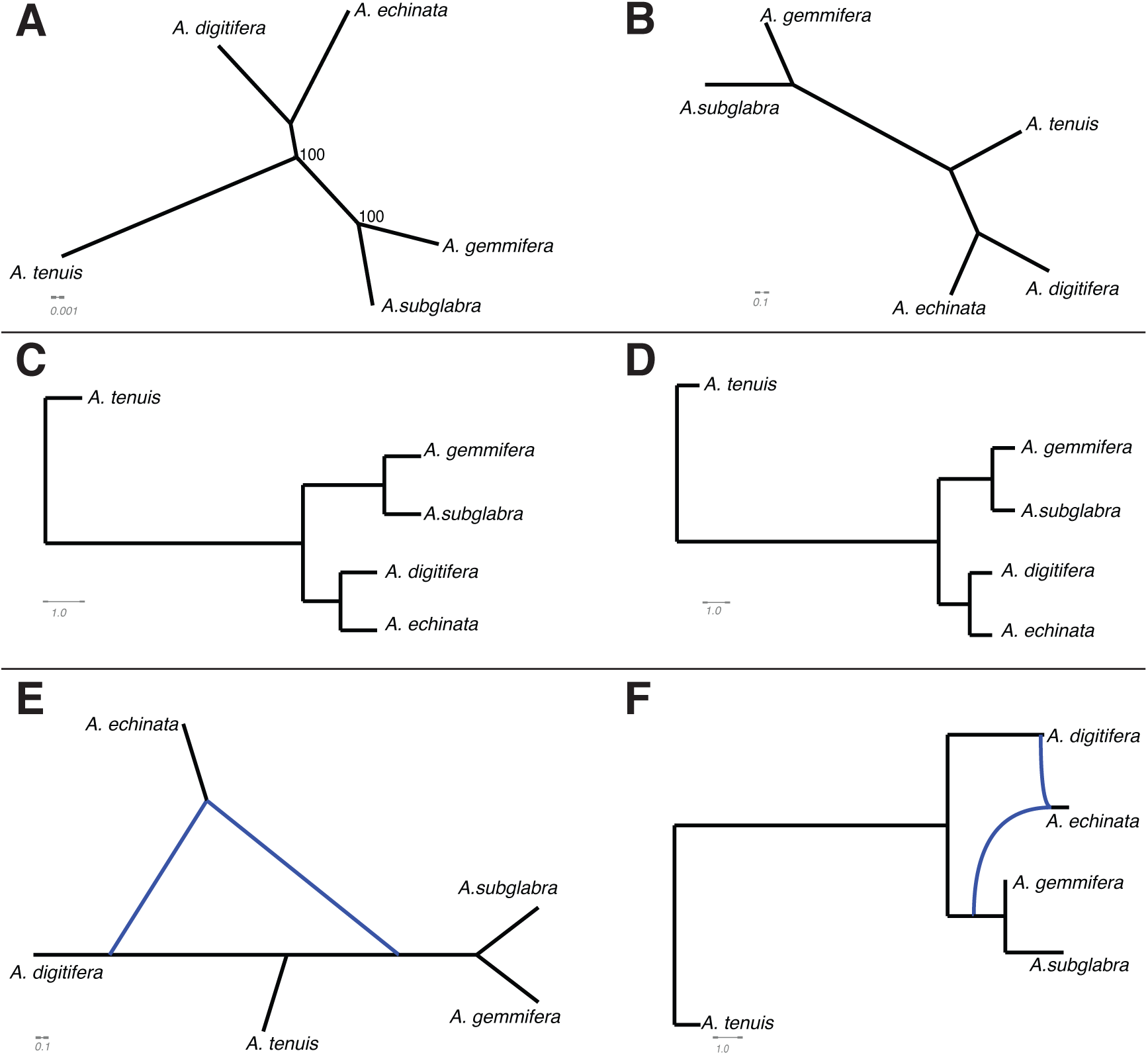
Validation of TREEasy on empirical data. TREEasy running on 4, 945 single-copy orthologs of *Acropora* generated species trees by (A) concatenated method, (B) ASTRAL, (C) MP-EST, and (D) STELLS2; and phylogenetic networks by (E) SNaQ and (F) PhyloNet.

### Inferring species trees and phylogenetic networks of the WGD clade of yeast

As a third test, we applied TREEasy to a novel analysis by analyzing data from a previous study that did not perform all the same analyses. The previous study investigated the evolutionary relationships of subphylum *Saccharomycotina* based on hundreds of yeast genomes (Shen et al., 2018). In particular, there is a clade (WGD clade) including common and important yeasts such as the baker’s yeast (Gonçalves et al., 2016; Ludlow et al., 2016), and there are two “non-robust internodes” in this clade. In addition, introgression has been reported in yeast (Leducq et al., 2016; Marcet-Houben and Gabaldón, 2015). Therefore, in order to investigate whether the “non-robust internodes” were caused by introgression, we first extracted sequences for 40 species from the WGD clade and an outgroup species (*Neurospora crassa*) with no missing data from two datasets (2408OG dataset and 1292BUSCO dataset). All horizontal gene transfer (HGT) genes were removed in these two datasets. 320 genes and 777 genes were extracted from 1292BUSCO and 2408OG datasets respectively and we applied TREEasy on these two datasets. Then, we found that the species trees inferred from different methods or datasets are not identical, and most of incongruences between the inferred species trees were located on the “non-robust internodes” (Figure 5).

**Figure 5.**
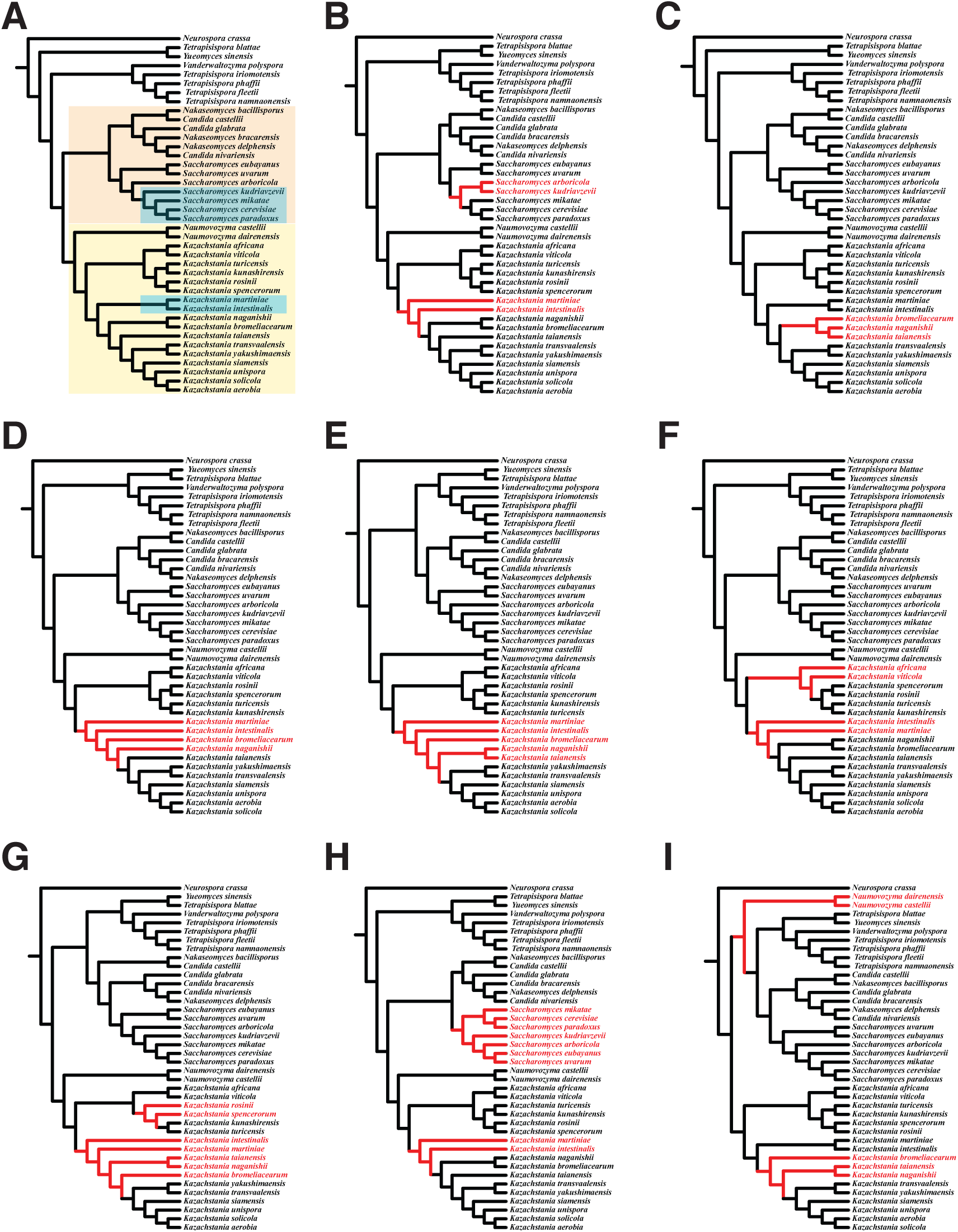
A case study of TREEasy on WGD clade of yeast genomic data. (A) The topology of WGD clade of yeast in the previous study. TREEasy running on two datasets generated species trees by (B, C) concatenated method, (D, E) ASTRAL, (F, G) MP-EST, and (H, I) STELLS2. The results (B, D, F, H) were generated by 1292BUSCO dataset and the results (C, E, G, I) were generated by 2408OG dataset. The blue shadows in (A) represents the “non-robust internodes” and the orange and yellow shadows represent two sub-clades: “*Saccharomyces*” clade and “*Kazachstania*” clade, for phylogenetic network analysis.

Next, considering running time of inferring phylogenetic network, we extracted the two sub-clades including “non-robust internodes” (“*Saccharomyces*” clade and “*Kazachstania*” clade) with two species as outgroups (*Yueomyces sinensis* and *Tetrapisispora blattae*) and run these two clades on TREEasy. After filtering the gene trees with bootstrap values smaller than 30 (B parameter), in 1292BUSCO dataset, we found 279 gene trees and 115 gene trees in “*Saccharomyces*” clade and “*Kazachstania*” clade, respectively. Meanwhile, in 2408BUSCO dataset, we found 654 gene trees and 351 gene trees in “*Saccharomyces*” clade and “*Kazachstania*” clade, respectively.

In order to reduce bias of species tree and phylogenetic network inferences from small gene tree numbers, we only reported the phylogenetic networks for “*Saccharomyces*” clade (Figure 6A) and “*Kazachstania*” clade (Figure 6B) in 2408BUSCO dataset here. On the one hand, we found some reticulate events occurred in the “non-robust internodes” in both clades. On the other hand, reticulate events occurred in some lineages which had the same topology from different species tree inference methods.

**Figure 6.**
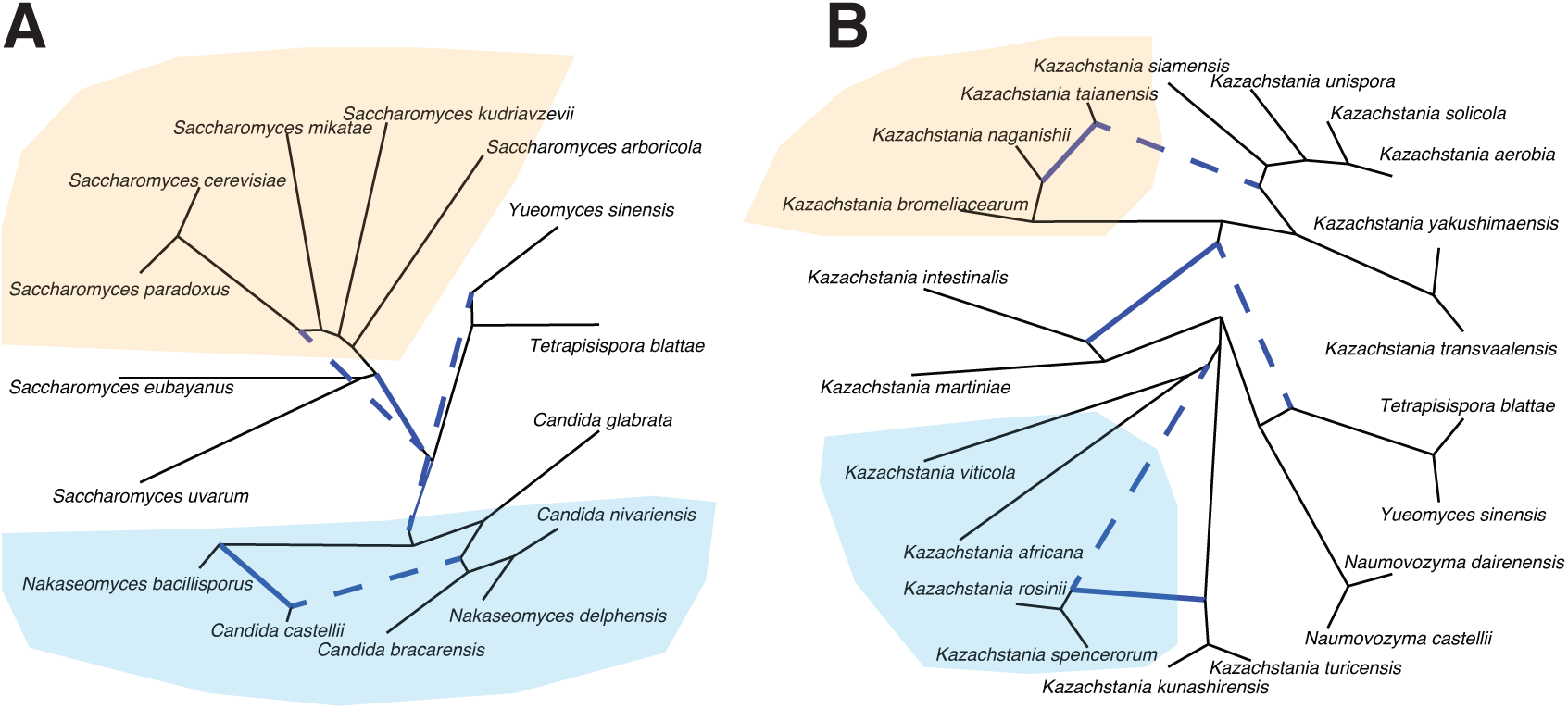
Phylogenetic network inferences for two sub-clades of yeast genomic data. Phylogenetic networks inferred by SNaQ on “*Saccharomyces*” clade (A) and “*Kazachstania*” clade (B) from 2408OG dataset. The orange shades represent reticulate events occurred in “non-robust internodes”. The blue shades represent reticulate events occurred in lineages which had the same topology from different species tree inference methods.

## Discussion

The era of big data and massive computing resources allow us to better understand species and population relationships (Allen et al., 2019; Bravo et al., 2019; Delsuc et al., 2005; Liu et al., 2015a). It is now possible to infer species trees and phylogenetic networks from hundreds of gene trees rather than concatenating a few loci to reconstruct a phylogeny. We developed a reliable and efficient tool called TREEasy to infer species trees and networks from molecular sequences directly. TREEasy is written in Python and can be run with multi-processors (https://github.com/MaoYafei/TREEasy).

The running time improved 6.5 times with 12 processors compared to with 1 processor, one of possible reasons is that parallelization by the threading module in python is applied to sequence alignment and gene tree reconstruction in TREEasy. Meanwhile, with gene or taxon number increasing, both running time and maximum memory usage increased as expected. Interestingly, the increase of taxon number has more effects on running time compared to the increase of gene number while the increase of gene number has more effects on memory usage compared to the increase of taxon number. One possible reason is that tree space searching results in increase of running time while heavy computation on gene tree reconstruction during parallelization leads to more memory requirements. In particular, phylogenetic network inference requires lots of running time because it is still difficult to infer phylogenetic network with a large taxon number (Solís-Lemus et al., 2017; Wen et al., 2018).

The empirical validation of five *Acropora* species generated the same result as our previous study (Mao et al., 2018), and thus, this result suggests TREEasy is a reliable tool for species tree and network inferences. Next, in order to evaluate TREEasy with more taxa, we applied TREEasy to a newly-sequenced yeast genomic data (Shen et al., 2018). Our results both confirm previous results, but go well beyond. First, the incongruence between the inferred species trees showed on two clades (“*Saccharomyces*” clade and “*Kazachstania*” clade) identical to the previous study. Moreover, the inferred phylogenetic networks suggest that the incongruence between the inferred species trees probably is caused by introgression. Interestingly, we also found some reticulate events occurred in the lineages which had the same topology from different species tree inferences. These results show that TREEasy would be easily applied to a genomic study and provide comprehensive insights on species relationships. It is worthy to note that each gene/locus has to contain all species sequences, namely, no missing data are allowed in TREEasy.

In all, this study presents a reliable and user-friendly tool to infer species tree and network from molecular sequences directly, potential to be used widely in population genetic/genomic, phylogenomic and phylogeographic studies. We hope that this will lower barriers to more comprehensive analyses of evolutionary history and accelerate research in systematic biology.

## ACKNOWLEDGEMENTS

This study was supported by OIST and was funded by JSPS grants (No.17J00557 to Y.M and Kakenhi No.17K15180 to E.P.E.). The authors acknowledge support of the OIST Scientific Computing Support Section. We additionally thank Nitish Narula for helping to test the program.

## AUTHOR CONTRIBUTIONS

Y.M and E.E conceived the study. Y.M performed all analyses in this study. S.H built the GUI. Y.M and E.E wrote the initial manuscript and edited the final manuscript.

## DATA ACCESSIBILITY

The following information was supplied regarding data availability: GitHub: https://github.com/MaoYafei/TREEasy.

## REFERENCES

Abascal, F., Zardoya, R., and Telford, M.J. (2010). TranslatorX: multiple alignment of nucleotide sequences guided by amino acid translations. Nucleic acids research 38, W7–13.

Allen, J.M., Germain-Aubrey, C.C., Barve, N., Neubig, K.M., Majure, L.C., Laffan, S.W., Mishler, B.D., Owens, H.L., Smith, S.A., and Whitten, W.M. (2019). Spatial Phylogenetics of Florida Vascular Plants: The Effects of Calibration and Uncertainty on Diversity Estimates. iScience 11, 57–70.

Barraclough, T.G., and Nee, S. (2001). Phylogenetics and speciation. Trends in ecology & evolution 16, 391–399.

Bastide, P., Solis-Lemus, C., Kriebel, R., William Sparks, K., and Ané, C. (2018). Phylogenetic comparative methods on phylogenetic networks with reticulations. Systematic biology 67, 800–820.

Berner, D., and Salzburger, W. (2015). The genomics of organismal diversification illuminated by adaptive radiations. Trends in genetics: TIG 31, 491–499.

Borowiec, M.L. (2016). AMAS: a fast tool for alignment manipulation and computing of summary statistics. PeerJ 4, e1660.

Bravo, G.A., Antonelli, A., Bacon, C.D., Bartoszek, K., Blom, M.P.K., Huynh, S., Jones, G., Knowles, L.L., Lamichhaney, S., and Marcussen, T. (2019). Embracing heterogeneity: coalescing the Tree of Life and the future of phylogenomics. PeerJ 7, e6399.

de Oliveira Martins, L., and Posada, D. (2017). Species tree estimation from genome-wide data with Guenomu. In Bioinformatics (Springer), pp. 461–478.

Degnan, J.H., and Rosenberg, N.A. (2009). Gene tree discordance, phylogenetic inference and the multispecies coalescent. Trends in ecology & evolution 24, 332–340.

Delsuc, F., Brinkmann, H., and Philippe, H. (2005). Phylogenomics and the reconstruction of the tree of life. Nature Reviews Genetics 6, 361.

Edwards, S.V., Liu, L., and Pearl, D.K. (2007). High-resolution species trees without concatenation. Proceedings of the National Academy of Sciences 104, 5936–5941.

Gadagkar, S.R., Rosenberg, M.S., and Kumar, S. (2005). Inferring species phylogenies from multiple genes: concatenated sequence tree versus consensus gene tree. Journal of Experimental Zoology Part B: Molecular and Developmental Evolution 304, 64–74.

Gonçalves, M., Pontes, A., Almeida, P., Barbosa, R., Serra, M., Libkind, D., Hutzler, M., Gonçalves, P., and Sampaio, J.P. (2016). Distinct domestication trajectories in top-fermenting beer yeasts and wine yeasts. Current Biology 26, 2750–2761.

Huson, D.H., and Bryant, D. (2005). Application of phylogenetic networks in evolutionary studies. Molecular biology and evolution 23, 254–267.

Kubatko, L.S., and Degnan, J.H. (2007). Inconsistency of phylogenetic estimates from concatenated data under coalescence. Systematic biology 56, 17–24.

Lambert, S.M., Reeder, T.W., and Wiens, J.J. (2015). When do species-tree and concatenated estimates disagree? An empirical analysis with higher-level scincid lizard phylogeny. Mol Phylogenet Evol 82, 146–155.

Leducq, J.-B., Nielly-Thibault, L., Charron, G., Eberlein, C., Verta, J.-P., Samani, P., Sylvester, K., Hittinger, C.T., Bell, G., and Landry, C.R. (2016). Speciation driven by hybridization and chromosomal plasticity in a wild yeast. Nature Microbiology 1, 15003.

Liu, L., Wu, S., and Yu, L. (2015a). Coalescent methods for estimating species trees from phylogenomic data. Journal of Systematics and Evolution 53, 380–390.

Liu, L., Xi, Z., Wu, S., Davis, C.C., and Edwards, S.V. (2015b). Estimating phylogenetic trees from genome-scale data. Annals of the New York Academy of Sciences 1360, 36–53.

Liu, L., and Yu, L. (2011). Estimating species trees from unrooted gene trees. Systematic biology 60, 661–667.

Liu, L., Yu, L., and Edwards, S.V. (2010). A maximum pseudo-likelihood approach for estimating species trees under the coalescent model. Bmc Evol Biol 10, 302.

Ludlow, C.L., Cromie, G.A., Garmendia-Torres, C., Sirr, A., Hays, M., Field, C., Jeffery, E.W., Fay, J.C., and Dudley, A.M. (2016). Independent origins of yeast associated with coffee and cacao fermentation. Current Biology 26, 965–971.

Mao, Y., Economo, E.P., and Satoh, N. (2018). The Roles of Introgression and Climate Change in the Rise to Dominance of Acropora Corals. Current Biology 28, 3373–3382. e3375.

Marcet-Houben, M., and Gabaldón, T. (2015). Beyond the whole-genome duplication: phylogenetic evidence for an ancient interspecies hybridization in the baker’s yeast lineage. PLoS biology 13, e1002220.

Mirarab, S., Reaz, R., Bayzid, M.S., Zimmermann, T., Swenson, M.S., and Warnow, T. (2014). ASTRAL: genome-scale coalescent-based species tree estimation. Bioinformatics 30, i541–i548.

Morrison, D.A. (2014). Is the tree of life the best metaphor, model, or heuristic for phylogenetics? Systematic biology 63, 628–638.

Nakamura, T., Yamada, K.D., Tomii, K., and Katoh, K. (2018). Parallelization of MAFFT for large-scale multiple sequence alignments. Bioinformatics 34, 2490– 2492.

Nguyen, L.-T., Schmidt, H.A., von Haeseler, A., and Minh, B.Q. (2014). IQ-TREE: a fast and effective stochastic algorithm for estimating maximum-likelihood phylogenies. Molecular biology and evolution 32, 268–274.

Page, R.D.M., and Charleston, M.A. (1997). From gene to organismal phylogeny: reconciled trees and the gene tree/species tree problem. Mol Phylogenet Evol 7, 231–240.

Pei, J., and Wu, Y. (2017). STELLS2: fast and accurate coalescent-based maximum likelihood inference of species trees from gene tree topologies. Bioinformatics 33, 1789–1797.

Shaw, T.I., Ruan, Z., Glenn, T.C., and Liu, L. (2013). STRAW: species TRee analysis web server. Nucleic acids research 41, W238–W241.

Shen, X.-X., Opulente, D.A., Kominek, J., Zhou, X., Steenwyk, J.L., Buh, K.V., Haase, M.A.B., Wisecaver, J.H., Wang, M., and Doering, D.T. (2018). Tempo and Mode of Genome Evolution in the Budding Yeast Subphylum. Cell 175, 1533-1545. e1520.

Shen, X.-X., Salichos, L., and Rokas, A. (2016). A genome-scale investigation of how sequence, function, and tree-based gene properties influence phylogenetic inference. Genome Biol Evol 8, 2565–2580.

Solís-Lemus, C., Bastide, P., and Ané, C. (2017). PhyloNetworks: a package for phylogenetic networks. Molecular biology and evolution 34, 3292–3298.

Soltis, D., and Soltis, P. (2018). The Great Tree of Life. (Academic Press).

Tonini, J., Moore, A., Stern, D., Shcheglovitova, M., and Ortí, G. (2015). Concatenation and species tree methods exhibit statistically indistinguishable accuracy under a range of simulated conditions. PLoS currents 7.

Wen, D., Yu, Y., Zhu, J., and Nakhleh, L. (2018). Inferring phylogenetic networks using PhyloNet. Systematic biology 67, 735–740.

Xu, S. (2000). Phylogenetic analysis under reticulate evolution. Molecular biology and evolution 17, 897–907.

